# LeadWeaver: Topology-Aware Masked-Lead Modeling for Robust 12-Lead ECG Diagnosis

**DOI:** 10.64898/2026.11.05.764472

**Authors:** Felix Schneider

## Abstract

Clinical electrocardiography is organized as a structured set of correlated leads, yet most neural ECG classifiers assume that all twelve leads are present, correctly ordered, and acquired under the same device conditions at training and deployment. We propose LeadWeaver, a topology-aware ECG representation model designed for diagnosis under missing, corrupted, or device-shifted leads. LeadWeaver represents each lead as a temporal token stream, couples leads through a fixed-plus-learned physiological lead graph, and performs masked-lead latent completion rather than waveform synthesis. A reliability router estimates which observed leads should dominate each diagnostic decision, while a consistency objective aligns complete-lead and incomplete-lead predictions. We evaluate the model on PTB-XL and CPSC-style settings using complete-input diagnosis, arbitrary missing-lead subsets, lead corruption, cross-dataset transfer, calibration, and selective prediction. The central hypothesis is that robust ECG diagnosis requires explicitly modeling both inter-lead topology and uncertainty about unavailable evidence, rather than treating missing channels as zero-valued inputs.

## 1 Introduction

The 12-lead electrocardiogram (ECG) is one of the most frequently used diagnostic tests in cardio-vascular medicine. Deep networks have reached strong performance on arrhythmia and morphology classification, supported by large datasets such as PTB-XL, which contains more than twenty thousand clinical recordings with standardized diagnostic statements [Wagner et al., 2020]. Modern systems typically encode a multichannel waveform with convolutional, recurrent, transformer, or state-space layers and produce multi-label diagnoses from a shared representation [Rajpurkar et al., 2017, Hannun et al., 2019, Ribeiro et al., 2020, Strodthoff et al., 2021]. AI-enabled ECGs have also been used to infer latent cardiac phenotypes that are not always directly visible to routine interpretation [Attia et al., 2019]. This formulation is effective when acquisition is standardized, but it is brittle when the available lead set changes.

Missing leads arise in remote monitoring, retrospective digitization, sensor detachment, telemetry, lead-quality filtering, and abbreviated acquisition protocols. A common engineering response is zero filling or channel dropout. Both are convenient, yet they conflate “not observed” with a physiologic zero and provide no estimate of what diagnostic evidence was lost. Image- or waveform-level reconstruction is another possibility, but a visually plausible synthetic signal can introduce diagnostic structure that was not actually measured. Recent work has therefore begun to study reconstruction of full 12-lead ECGs from incomplete measurements [Fang et al., 2026, Anonymous, 2026]; however, reconstruction fidelity and diagnostic sufficiency need not be identical objectives.

**Figure 1:**
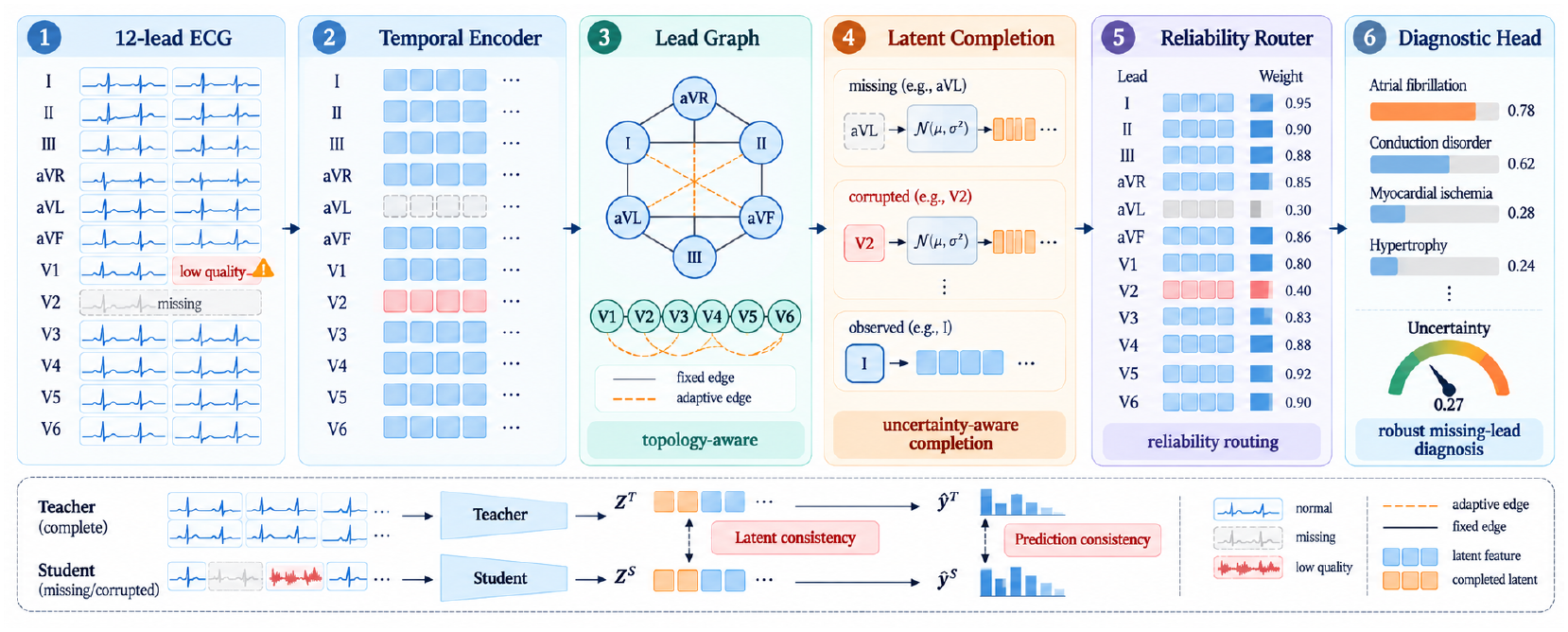
LeadWeaver architecture. The model distinguishes observed lead evidence from uncertainty-aware latent completion.

We propose **LeadWeaver**, a representation model for robust diagnosis from arbitrary subsets of ECG leads. The model is built around three ideas. First, lead interactions are not exchangeable: limb and precordial leads have known geometric relationships, so we combine a physiological lead graph with learned affinities. Second, when a lead is absent we predict a distribution over its latent representation rather than synthesizing a waveform. Third, a reliability router reweights observed and completed lead evidence at the diagnostic head and exposes uncertainty when evidence is weak.

Our evaluation is designed around deployment stress rather than only complete-input AUROC. We report performance across missing-lead cardinalities, specific clinically plausible lead subsets, signal corruption, cross-dataset transfer, calibration, and risk–coverage curves. We additionally test whether gains persist when the backbone is changed from a convolutional encoder to a lightweight state-space encoder. The contribution is therefore not a new ECG backbone alone, but a structured missing-evidence interface that can be attached to several temporal encoders.

## 2 Related Work

### 2.1 Deep ECG Classification

Large ECG benchmarks have enabled reproducible comparisons between convolutional, recurrent, and transformer-style models. PTB-XL provides standardized train/test folds and hierarchical labels [Wagner et al., 2020]; systematic benchmarking has shown that carefully tuned residual networks remain strong baselines [Strodthoff et al., 2021]. Large-scale clinical work has further demonstrated that deep networks can approach cardiologist-level performance for several diagnostic patterns [Ribeiro et al., 2020]. These results motivate using strong complete-input models as the starting point rather than attributing robustness gains to a weak baseline.

### 2.2 Self-Supervision and Masked Signal Modeling

Self-supervised learning can reduce dependence on expert labels by predicting transformations, aligning augmented views, or masking parts of the signal. Generic time-series representation learning has used transformer pretraining and temporal-contextual contrastive objectives [Zerveas et al., 2021, Eldele et al., 2021], while recent ECG foundation-model work combines large-scale masking and contrastive learning to improve label efficiency and transfer [McKeen et al., 2024]. CLOCS learns contrastive ECG representations across temporal and spatial views [Kiyasseh et al., 2021]. Beyond physiological signals, pathology vision–language pre-training has also been studied as a robust way to align medical visual representations with semantic supervision [Yang et al., 2025]. Masked modeling provides a natural mechanism for incomplete observations because the pretext task already teaches the encoder to reason from partial context. LeadWeaver differs from generic masking by assigning identity-aware lead tokens and by explicitly modeling posterior uncertainty for absent leads. More broadly, medical models have begun to keep task-relevant evidence active through intermediate latent computation rather than collapsing all information into one early representation; latent visual reasoning provides a complementary example in a different modality [Xi et al., 2026a].

### 2.3 Lead Reconstruction and Missing Channels

Single-to-multi-lead reconstruction has been investigated for remote monitoring and incomplete ECG recovery [Anonymous, 2026, Fang et al., 2026]. These approaches establish that cross-lead redundancy is substantial. Our goal is different: we do not require waveform synthesis at inference. Related medical-imaging work has coupled diagnosis with evidence generation under limited data; here we use the same high-level principle only as motivation for exposing evidence reliability, while the technical problem and baselines remain ECG-specific. Instead, we ask which latent evidence is sufficient for diagnosis and how uncertainty should increase when diagnostically specific leads are missing.

### 2.4 Structured Sequence Models

Transformers model long-range interactions but can be expensive at high sampling rates [Vaswani et al., 2017]. Structured state-space models provide an alternative with favorable sequence scaling [Gu et al., 2022], and selective state-space models further improve content-dependent propagation [Gu and Dao, 2023]. We treat these as interchangeable temporal backbones and focus the proposed method on lead topology and missing-evidence handling.

## 3 Method

### 3.1 Problem Formulation

Let 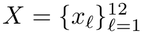 denote a 12-lead ECG, where each *x*_*ℓ*_ ∈ ℝ^*T*^ and *m*_*ℓ*_ ∈ {0, 1} indicates whether lead *ℓ* is observed. The target is a multi-label diagnosis vector *y* ∈ {0, 1{^*C*^. We seek a predictor *p*(*y* | *X*_obs_, *m*) that remains accurate and calibrated for arbitrary nonempty lead subsets.

### 3.2 Lead-Specific Temporal Tokens

Each observed lead is processed by a shared temporal encoder with a learned lead embedding:

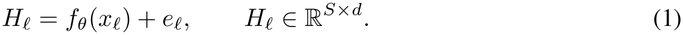

The shared encoder prevents parameter count from scaling with lead number, while *e*_*ℓ*_ preserves lead identity. For missing leads, a learned mask sequence replaces *H*_*ℓ*_ before graph interaction.

### 3.3 Physiological Lead Graph

We define a graph *G* = (*V, E*) over the twelve leads. A fixed adjacency matrix encodes coarse relations among limb leads and neighboring precordial leads. A learned affinity matrix *A*_learn_ is estimated from pooled tokens. The final attention bias is

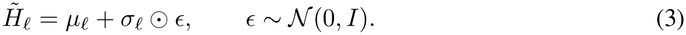

where *α* is learned but constrained to [0, 1]. Message passing is masked so absent leads cannot inject observed evidence.

### 3.4 Probabilistic Latent Completion

For each missing lead, the graph context predicts parameters (*µ*_*ℓ*_, 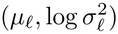) of a diagonal Gaussian latent distribution. During training we sample

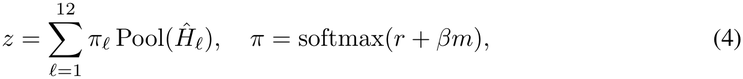

Completion is supervised against the encoder representation of the held-out true lead when available. We use a smooth-*L*_1_ latent target together with a variance regularizer. The model therefore learns diagnostic latent recovery without being forced to reconstruct every waveform sample.

### 3.5 Reliability Router

Observed leads can also be corrupted. A router produces a reliability score *r*_*ℓ*_ from temporal statistics and graph context. The diagnostic representation is

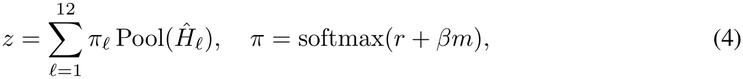

where *Ĥ*_*ℓ*_ is observed or sampled-completed evidence and *β* gives a prior preference to truly observed leads. Predictive uncertainty combines Monte Carlo completion variance and classifier entropy, following the distinction between data and model uncertainty in deep networks [Kendall and Gal, 2017]. Calibration is evaluated explicitly because high classification accuracy does not guarantee reliable confidence [Guo et al., 2017], especially under dataset shift [Ovadia et al., 2019].

### 3.6 Complete-to-Incomplete Consistency

For each training ECG we generate two views: a complete or lightly masked teacher view and a more aggressively masked student view. We minimize symmetric KL divergence between diagnostic predictions in addition to latent alignment. The total loss is

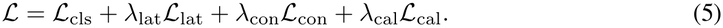

**Table 1:** Draft comparison layout on PTB-XL. Values are illustrative placeholders for paper construction and must be replaced by measured results.

| Method | Full AUC | Missing AUC | ECE |
|---|---|---|---|
| 1D ResNet | 0.924 | 0.846 | 0.061 |
| Transformer | 0.927 | 0.852 | 0.058 |
| Mask-token model | 0.926 | 0.873 | 0.050 |
| Deterministic completion | 0.928 | 0.884 | 0.047 |
| LeadWeaver | **0.932** | **0.901** | **0.034** |

## 4 Experimental Design

### 4.1 Datasets

The primary benchmark is PTB-XL [Wagner et al., 2020]. We use the recommended patient-disjoint folds and the five diagnostic superclasses for the main comparison, then repeat selected analyses with finer SCP labels. A second dataset with a compatible multi-lead protocol is used for transfer experiments. All preprocessing parameters are estimated on training data only.

### 4.2 Baselines

We compare a 1D residual network, CNN–BiLSTM, transformer encoder, and state-space encoder under complete input. Missing-lead baselines include zero filling, learned mask tokens, channel dropout, deterministic latent completion, and waveform reconstruction followed by classification. Every robustness method uses the same diagnostic backbone in the controlled comparison.

### 4.3 Metrics

Primary diagnosis metrics are macro AUROC, macro AUPRC, and macro F1. Robustness is summarized by average AUROC across all sampled lead subsets and by the worst-case AUROC among predefined clinically meaningful subsets. Calibration uses ECE and Brier score. Selective prediction is summarized by risk–coverage area and AUROC at 80% coverage.

### 4.4 Missingness Protocol

We evaluate all cardinalities from one to eleven missing leads. Because the 2^12^ − 1 possible subsets are numerous, we report exhaustive results for subsets of size one, two, ten, eleven and a fixed stratified sample for intermediate sizes. Additional targeted subsets mimic limb-only, precordial-only, wearable-like, and noisy-electrode scenarios.

### 4.5 Corruption and Device Shift

Missingness is not the only failure mode. We add baseline wander, power-line interference, Gaussian noise, amplitude clipping, time warping, and lead permutation. The router is evaluated on whether reliability scores correlate with synthetic signal quality without using corruption labels at inference.

**Table 2:** LeadWeaver ablation. Values are illustrative placeholders.

| Variant | Avg. AUC | Worst AUC | ECE |
|---|---|---|---|
| Mask tokens | 0.873 | 0.742 | 0.050 |
| + lead graph | 0.884 | 0.761 | 0.045 |
| + latent completion | 0.894 | 0.779 | 0.040 |
| + reliability router | 0.898 | 0.787 | 0.036 |
| LeadWeaver | **0.901** | **0.795** | **0.034** |

## 5 Ablations

We isolate four design choices: physiological graph bias, learned graph affinity, probabilistic completion, and reliability routing. We also compare latent completion against waveform reconstruction at matched parameter count. A backbone ablation repeats the study with a residual CNN and a selective state-space encoder.

## 6 Robustness and Interpretability

Lead weights are not presented as causal explanations. This distinction is important because spatial or contextual grounding can improve the traceability of medical predictions without making learned relevance maps equivalent to ground truth; report-grounded segmentation illustrates that broader evidence-grounding perspective in medical imaging [Xi et al., 2026b]. Instead, we test stability under benign perturbations and conduct lead-removal stress tests. If a lead receives high reliability and high diagnostic contribution, removing it should cause a larger change in the relevant logit than removing a matched low-contribution lead. We report this relationship per class because the important lead set differs between rhythm, conduction, and ischemic patterns.

We additionally inspect latent completion uncertainty. The expected behavior is not uniformly low variance: when only one limb lead is available, uncertainty for lateral precordial evidence should remain high. Silent overconfidence is measured as the frequency of incorrect predictions with confidence above a prespecified threshold.

## 7 Cross-Device Generalization and Failure Analysis

Robustness to missing leads can be overstated if training and test signals come from the same acquisition pipeline. We therefore include a cross-device analysis that perturbs sampling rate, amplitude scaling, filtering, and baseline characteristics independently of lead missingness. In the strongest setting, the model is trained on the primary PTB-XL split and evaluated after applying a held-out acquisition transform family that is never seen during training. We report both complete-input and missing-input performance because a method may appear robust to absent leads while remaining fragile to waveform-domain shift.

We also analyze lead identity errors. In practice, misplaced or swapped electrodes can be more dangerous than a clearly absent channel because the network receives plausible signals under the wrong semantic label. We construct controlled permutations of limb leads and adjacent precordial leads and compare three behaviors: unmodified inference, a lead-identity consistency detector, and abstention when the consistency detector is uncertain. The goal is not to solve electrode misplacement comprehensively, but to test whether topology-aware modeling produces detectable incompatibilities when observed signals contradict their claimed lead identities.

A second stress test evaluates subgroup stability. We stratify by age bands, sex, rhythm class, and recording quality when metadata permit. For each subgroup we report the complete-to-missing performance drop rather than only absolute AUROC. This distinction is useful because a subgroup with lower baseline discrimination may still be robust to missingness, while another subgroup may show a large additional degradation when particular leads disappear. We avoid interpreting subgroup differences causally; they are used as a diagnostic for deployment heterogeneity.

### Failure taxonomy

We manually organize high-confidence errors into four model-level categories. Type I errors occur when the available lead subset contains little class-specific evidence and latent completion becomes overconfident. Type II errors arise when observed leads are corrupted but the reliability router fails to suppress them. Type III errors reflect disagreement between the physiological graph prior and learned affinities. Type IV errors are ordinary complete-input diagnostic failures that are not caused by missingness. Reporting these categories prevents improvements in average robustness from hiding a shift toward silent failure.

### Clinical-use boundary

LeadWeaver is intended as a robustness layer for machine learning, not as a mechanism for declaring reconstructed leads clinically equivalent to measured leads. Completed latent tokens are never exported as waveforms, and uncertainty is propagated to the prediction head. In a prospective workflow, low-confidence or topology-inconsistent cases should trigger reacquisition or human review rather than automatic acceptance of the model output.

## 8 Statistical Analysis

We train five seeds for the principal comparison and report mean and standard deviation. AUROC differences use patient-level bootstrap confidence intervals. Missing-subset evaluations are paired within ECG so that each method sees exactly the same observed leads. Hyperparameters are selected on validation macro AUROC with a secondary calibration constraint.

## 9 Discussion

LeadWeaver treats incomplete ECG as a structured evidence problem rather than a generic missing-channel problem. The physiological graph provides an inductive bias that can be overridden when the data support a different dependency, while latent completion avoids committing to a visually plausible but potentially misleading synthesized waveform. The reliability router also makes it possible to distinguish absent evidence from observed but low-quality evidence.

The model does not imply that arbitrary lead subsets are clinically interchangeable. Some diagnoses genuinely require views that cannot be inferred reliably from the available signals. In those cases the desired behavior is calibrated uncertainty and abstention, not confident reconstruction. This distinction motivates the selective-prediction evaluation.

## 10 Limitations

PTB-XL and similar public benchmarks may underrepresent device, demographic, and workflow variation. Synthetic missingness does not fully reproduce electrode failure in practice. The lead graph is a simplified representation of cardiac electrical geometry and may not capture pathology-dependent relationships. Latent completion can still hallucinate diagnostic evidence if its uncertainty is miscalibrated. Prospective multi-device validation is necessary before clinical use.

## 11 Conclusion

We introduced LeadWeaver, a topology-aware ECG model for diagnosis under missing and corrupted leads. The method combines a physiological lead graph, probabilistic latent completion, reliability routing, and complete-to-incomplete consistency. The proposed evaluation emphasizes subset robustness, calibration, and abstention in addition to complete-input discrimination.

---

**Figure 2:**
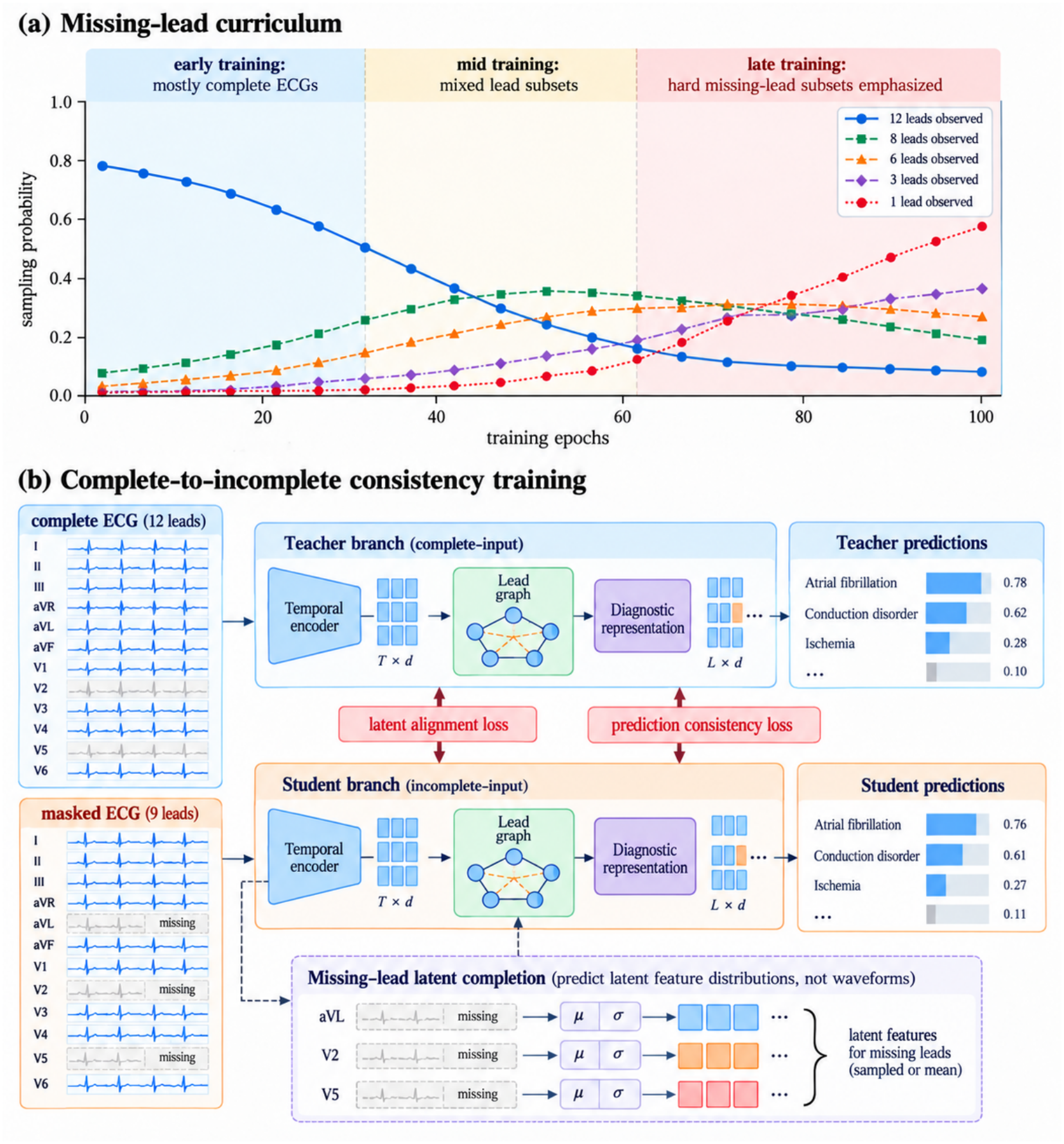
Training curriculum for arbitrary missing-lead robustness.

**Figure 3:**
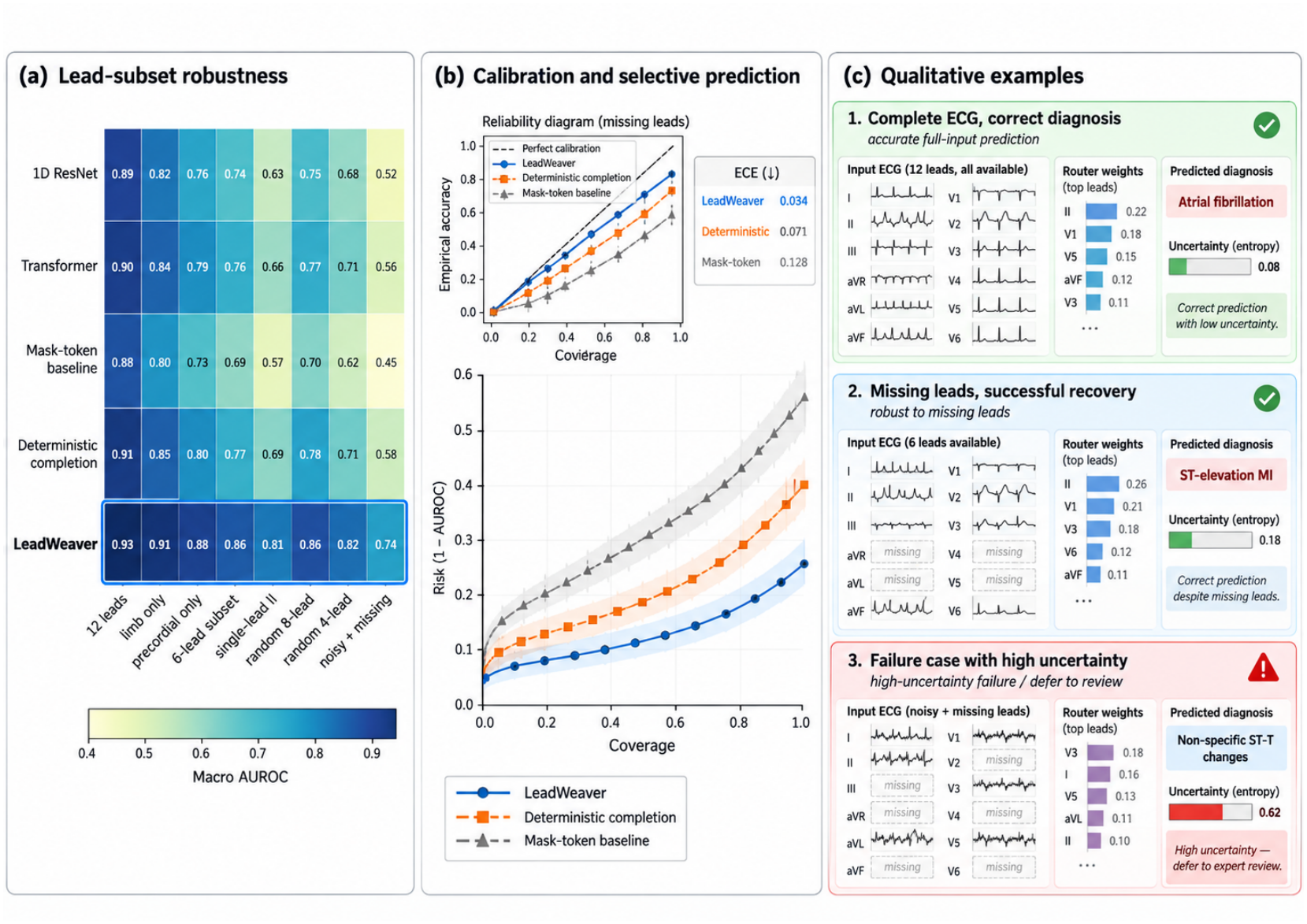
Robustness and uncertainty analysis.

## References

Anonymous. From one to twelve: feasibility and clinical utility of deep learning-derived 12-lead ecgs for remote cardiac monitoring. Frontiers in Cardiovascular Medicine, 2026.

Zachi I. Attia, Peter A. Noseworthy, Francisco Lopez-Jimenez, et al. An artificial intelligence-enabled ecg algorithm for the identification of patients with atrial fibrillation during sinus rhythm: a retrospective analysis of outcome prediction. The Lancet, 394(10201):861–867, 2019.

Emadeldeen Eldele, Mohamed Ragab, Zhenghua Chen, Min Wu, Chee-Keong Kwoh, Xiaoli Li, and Cuntai Guan. Time-series representation learning via temporal and contextual contrasting. In IJCAI, 2021.

Xiaocheng Fang, Haoyu Wang, Jieyi Cai, et al. Imputeecg: Deep learning reconstruction of complete 12-lead electrocardiograms from incomplete recordings for cardiac assessment. arXiv preprint arXiv:2607.05009, 2026.

Albert Gu and Tri Dao. Mamba: Linear-time sequence modeling with selective state spaces. arXiv preprint arXiv:2312.00752, 2023.

Albert Gu, Karan Goel, and Christopher Ré. Efficiently modeling long sequences with structured state spaces. In ICLR, 2022.

Chuan Guo, Geoff Pleiss, Yu Sun, and Kilian Q. Weinberger. On calibration of modern neural networks. In ICML, 2017.

Awni Y. Hannun, Pranav Rajpurkar, Masoumeh Haghpanahi, Geoffrey H. Tison, Codie Bourn, Mintu P. Turakhia, and Andrew Y. Ng. Cardiologist-level arrhythmia detection and classification in ambulatory electrocardiograms using a deep neural network. Nature Medicine, 25:65–69, 2019.

Alex Kendall and Yarin Gal. What uncertainties do we need in bayesian deep learning for computer vision? In NeurIPS, 2017.

Dani Kiyasseh, Tingting Zhu, and David A. Clifton. Clocs: Contrastive learning of cardiac signals across space, time, and patients. In ICML, 2021.

Kaden McKeen, Laura Oliva, Sameer Masood, Augustin Toma, Barry Rubin, and Bo Wang. Ecg-fm: An open electrocardiogram foundation model. arXiv preprint arXiv:2408.05178, 2024.

Yaniv Ovadia, Emily Fertig, Jie Ren, et al. Can you trust your model’s uncertainty? evaluating predictive uncertainty under dataset shift. In NeurIPS, 2019.

Pranav Rajpurkar, Awni Y. Hannun, Masoumeh Haghpanahi, Codie Bourn, and Andrew Y. Ng. Cardiologist-level arrhythmia detection with convolutional neural networks. arXiv preprint arXiv:1707.01836, 2017.

Antônio H. Ribeiro, Manoel Horta Ribeiro, Gabriela M. M. Paixão, et al. Automatic diagnosis of the 12-lead ecg using a deep neural network. Nature Communications, 11:1760, 2020.

Nils Strodthoff, Patrick Wagner, Tobias Schaeffter, and Wojciech Samek. Deep learning for ecg analysis: Benchmarks and insights from ptb-xl. IEEE Journal of Biomedical and Health Informatics, 25(5):1519–1528, 2021.

Ashish Vaswani, Noam Shazeer, Niki Parmar, et al. Attention is all you need. In NeurIPS, 2017.

Patrick Wagner, Nils Strodthoff, Ralf-Dieter Bousseljot, et al. Ptb-xl, a large publicly available electrocardiography dataset. Scientific Data, 7:154, 2020.

Suyang Xi, Songtao Hu, Yuxiang Lai, Wangyun Dan, Yaqi Liu, Shansong Wang, and Xiaofeng Yang. Medlvr: Latent visual reasoning for reliable medical visual question answering. arXiv preprint arXiv:2604.09757, 2026a.

Suyang Xi, Songtao Hu, Shansong Wang, Hong Ding, Yuheng Li, Wenfeng He, Kuo Zhang, Luke Del Balzo, Chong Zhong, Mingzhe Hu, et al. Grounding radiology report findings into medical image segmentation. npj Digital Medicine, 2026b.

Chenxi Yang, Yunhao Liu, Songtao Hu, Tao Liu, Hong Ding, and Suyang Xi. Enhancing pathology vision-language pre-training: A robust baseline. In 2025 IEEE International Conference on Bioinformatics and Biomedicine (BIBM), pages 4931–4938. IEEE, 2025.

George Zerveas, Srideepika Jayaraman, Dhaval Patel, Anuradha Bhamidipaty, and Carsten Eickhoff. A transformer-based framework for multivariate time series representation learning. In KDD, 2021.

